# Enterprise Health Twins - Conceptual Framework, Design Principles, and Longitudinal Evaluation: Preliminary Results from EpiToMe

**DOI:** 10.64898/2025.12.06.692733

**Authors:** Guo-Qiang Zhang, Licong Cui, Wei-chun Chou, Johnson P. Hampson, Samden Lhatoo, Shiqiang Tao

## Abstract

Monolithic Electronic Health Record (EHR) architectures remain fundamentally misaligned with high-resolution, specialty-specific clinical workflows, contributing to significant physician burnout and challangies in secondary use of data. This study introduces the Enterprise Health Twin (EHT) framework, a paradigm grounded in the Digital Twin concept, with the goal to transform static clinical ledgers into dynamic decision-support systems. The EHT establishes continuous bidirectional feedback (feed-forward ingestion and feed-backward decision support) between the physical entities of clinical workflows and the digital counterparts of interfaces reflecting the evolving states of the workflow, for each individual patient in the EHT. We conceptualized, prototyped, and operationalized the EHT framework for epilepsy care for seven years, in production, at the point of care. Deployed at the Epilepsy Center of the University of Texas Health Science Center at Houston, a ∼400% increase productivity has been achieved in specialized surgical evaluation throughput without additional staffing. The EHT achieved de novo capture of ontology-anchored structured data at the point of care, mitigating cognitive load while growing the volume of high-resolution, annotated longitudinal datasets for clinical research, and driving an increase of care velocity in a learning healthcare system.

## 1 Introduction

The widespread digitization of medical data, catalyzed by the Health Information Technology for Economic and Clinical Health (HITECH) Act, successfully transitioned the United States healthcare system away from paper records^1^. However, the resulting enterprise Electronic Health Record (EHR) ecosystems have largely functioned as administrative requirements rather than cognitive augmentations. Current EHR architectures are fundamentally monolithic, designed primarily to optimize billing capture, coding compliance, and generic ambulatory triage. Consequently, they remain structurally misaligned with the longitudinal and high-resolution demands of modern, specialized clinical medicine^2^.

This misalignment is a well-documented driver of clinical inefficiency and the physician burnout epidemic. Time-motion informatics studies consistently indicate that physicians spend approximately two hours on clerical documentation and chart navigation for every hour of direct patient care, a burden that frequently extends into after-hours documentation^3^. This operational friction is acutely evident in multidisciplinary environments such as the Epilepsy Monitoring Unit (EMU). The presurgical evaluation of drug-resistant epilepsy requires clinicians to synthesize multimodal data streams over time, including continuous video-electroencephalography (vEEG), metabolic neuroimaging (FDG-PET, SPECT), structural morphometry, and invasive cortical mapping^7^. Standard EHRs relegate this multi-dimensional data into unstructured clinical notes scattered across disparate subsystems, failing to provide a cohesive picture of the patient’s longitudinal trajectory.

To resolve these systemic informatics deficiencies, we introduce a fundamental conceptual and architectural shift: the transition from passive, retrospective health records to dynamic **Enterprise Health Twins (EHTs)**. By adapting the principles of Digital Twins from engineering into clinical care^4,17,18,^ we represent and capture patient-care workflows that are dynamically reflected in their digital counterparts as live, interactive dashboards and interfaces (**Fig. 1**), leading to enhanced care pathway follow up and accelerated care delivery velocity.

**Figure 1.**
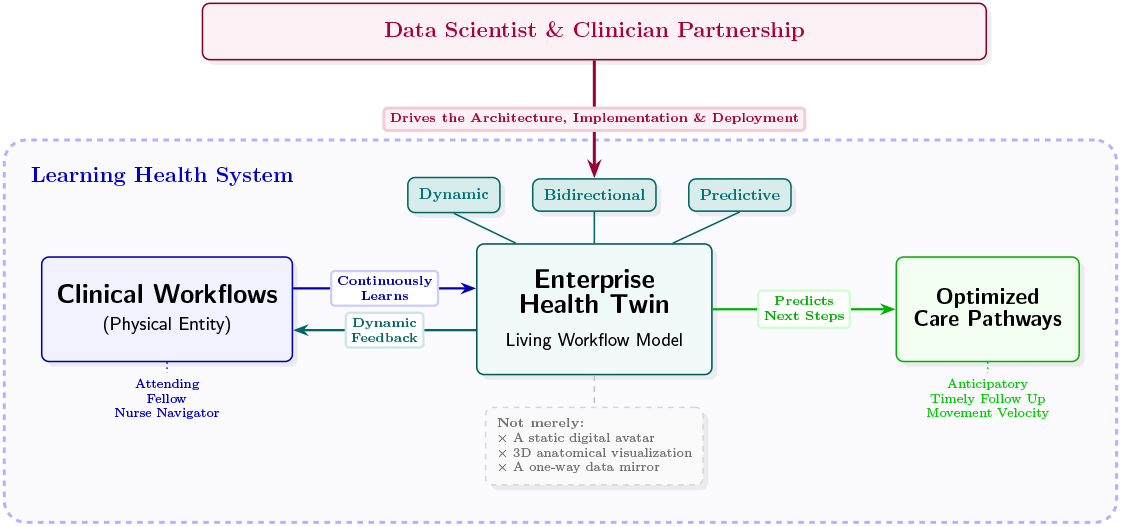
The central concept of EHT consisting of bidirectional feedback between clinical workflows and their digital representations (middle), leading to optimized care pathways.

**Figure 2.**
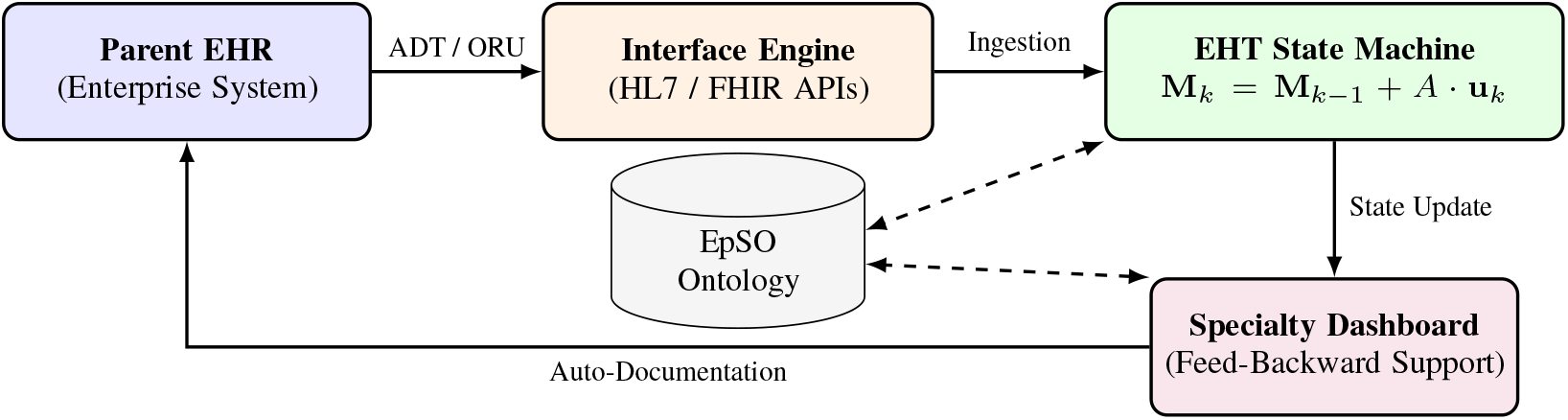
Architecture of the EpiToMe Enterprise Health Twin. The system bridges the enterprise EHR with a bespoke specialty dashboard through ontology-anchored feed-forward and feed-backward loops.

This paper elucidates the conceptual foundations of the EHT framework, formalizes its guiding principles for implementation, and provides an extensive longitudinal evidence of its impact in a Level 4 Comprehensive Epilepsy Center, a specialized medical facility that provides the highest level of medical and surgical evaluation and treatment for patients with complex, uncontrolled, or drug-resistant epilepsy.

## 2 Background

### Digital Twins

Based on the formal consensus definitions established by the U.S. National Academies of Sciences, Engineering, and Medicine (NASEM^5^), a true Digital Twin transcends a mere static digital model or one-way digital shadow. It must demonstrate automated, bidirectional, real-time data flow, predictive capabilities beyond available data, and the provision of actionable decision support to realize clinical value^18^. For example, a medical digital twin should be a living computational model that continuously learns from the patient, predicting outcomes from activities, nutrition, and pharmacological interventions to suggest optimized care pathways before adverse clinical events manifest. While recent biomedical digital twin research has predominantly focused on localized biophysical modeling (e.g., *in silico* physiological simulations^4^) our EHT framework uniquely expands this paradigm to encompass the *operational, cognitive, and clinical process layers* of healthcare delivery^18^ in a Learning Health System^6^ (**Fig. 1**).

Current EHRs operate reactively as “systems of record.” In contrast, EHT functions as an active cybernetic control system, sustained by continuous bidirectional information loops^5,18^: 1. *Feed-Forward Synchronization (Physical-to-Digital):* Contextual data from the care environment (e.g., HL7 Admission/Discharge/Transfer [ADT] messages, laboratory results, and imaging metadata) is continuously streamed to update the digital twin. This data acts as an event trigger that advances the patient within a predefined clinical state machine; 2. *Feed-Backward Support (Digital-to-Physical):* By evaluating the twin’s state against an underlying clinical care guideline, the system contextually surfaces specific clinical checklists, highlights missing diagnostic prerequisites, and automates the pre-population of specialty-specific documentation templates to lower cognitive load.

### EpiToMe: A Bespoke EHR for Epilepsy Care

We have developed EpiToMe (Epilepsy Tracking and optimization Management Engine^7^), which is a bespoke EHR system customized for epilepsy care. EpiToMe provides patient data capture functions for electroencephalogram (EEG) report, daily report and phase report for EMUs. It uses a domain-specific epilepsy and seizure ontology (EpSO) to: (1) support structured entry of multi-modal epilepsy data;(2) proactively ensure quality of data through use of ontology terms in nested facet systems; and (3) organize and index patient information for subsequent analytical queries and secondary use. EpiToMe has been developed following web-interface-driven development, an agile software development methodology, in close collaboration with physicians. EpiToMe has a built-in physician dashboard optimized for physicians to perform major tasks with “just-in-time” presentation of contextualized, action-ready interfaces. EpiToMe’s data entry supports collaborative editing: it allows all members of the care team, such as EEG technicians and clinical fellows, to take ownership of specific document subsections of their respective oversight. A 10-question survey was completed by 11 senior clinical attending physicians. Consensus was found that EpiToMe eased the burden of care documentation for patient management, a contributing factor to physician burnout^7^. In the past seven years, we have continuous expanded the clinical workflow scope of EpiToMe, documenting ∼185 thousand clinical reports for over 40 thousand unique patients.

## 3 Methods

The conceptual framework of Enterprise Health Twins proposed in this paper treats clinical workflows as a twining component rather than a passive record. The EHT is tethered to the enterprise EHR through health-data interfaces^2^. The EHT’s physical counterpart is the patient-care process involving attending physicians, fellows, technologists, nurse navigators, conferences, diagnostic tests, reports, and follow-up. The bidirectional loop is explicit: feed-forward synchronization moves clinical events such as admissions, reports, studies, imaging metadata, and documentation state, from the care environment into the digital counterpart. Feed-back support pushes context-aware checklists, status displays, missing-prerequisite alerts, and documentation assistance back into clinical work, and inform decision-making at the point of care. *EpiToMe is therefore an EHT*^*18*^.

### Colored Workflow Petri Nets

EHT’s capacity as a clinical workflow digital twin can be formulated mathematically as a *Colored Petri Net* ^*14–16*^. By adopting a mechanistic Petri Net framework, mathematical tokens carry rigorously typed, complex data payloads (e.g., HL7^®^ FHIR^®^ resources, and data capturing form completion status), enforcing care transition logic in clinical workflows.

A clinical EHT workflow be defined as a tuple ⟨*P, T, F*, Σ, *C*, Φ, **M**_0_⟩, where:

- *P* = { *p*_1_, …, *p*_*m*_} is a finite set of *places*, representing clinical prerequisites and real-time physiological states (e.g.,*p*_PET_ = “FDG-PET Completed”).
- *T* = {*t*_1_, …, *t*_*n*_} is a finite set of *transitions*, representing bidirectional clinical decision events and therapeutic interventions (e.g., *t*_Surg_ = “Proceed to Surgical Resection”).
- *F* ⊆ (*P* × *T*) ∪ (*T* × *P*) is the directed flow relation detailing the chronologically and causally permissible workflow mapping. For a transition *t*_*j*_ we write its *preset* (input places) and *postset* (output places) as ^*•*^*t*_*j*_ = { *p*_*i*_ ∈ *P* : (*p*_*i*_, *t*_*j*_) ∈ *F*} and 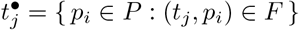.
- Σ is a finite set of *color sets*^15^. Each color set is a product *σ* = *τ* × Γ pairing (i) an ontology-anchored *semantic type τ* (an EpSO/LOINC/FHIR concept that guarantees data are structured rather than free text) with (ii) a totally ordered *lifecycle-status lattice* Γ that the deployed interfaces render as color. A token at a place is thus a pair ⟨*d, γ*⟩carrying a typed clinical datum *d*∈*τ* and a status *γ*∈ Γ.
- *C* : *P*→ Σ is the color function assigning to each place its interoperable schema *τ* and shared status lattice Γ.
- Φ : *T*× *M* → { 0, 1 } is the *predictive gating function*. Representing the twin’s continuous-learning capability, where Φ computationally evaluates the expected clinical utility and risk of a transition based on the patient’s current high-dimensional state within a “marking space” *M* .
- **M**_*k*_ ∈*M* is the colored multi-set marking vector at discrete temporal step *k*, representing the living, dynamically updated state of the EHT via continuous bidirectional patient-specific care input stream.

The dynamic state evolution is governed by the augmented state equation **M**_*k*_ = **M**_*k™*1_ + *A* · **u**_*k*_(*D*_in_) where *A* is the incidence matrix mapping transitions to places (evaluating over the semantic data types Σ), and **u**_*k*_(*D*_in_) is the control vector of triggered, standard-compliant data events dynamically driven by the patient care input stream (*D*_in_).

To prevent unquantified risks and safeguard the workflow structure, a clinical transition *t*_*j*_ is mathematically disabled unless two strict conditions hold: 1. *Semantic Process Conformance:* All upstream clinical prerequisites possess a valid, semantically mapped token (i.e., ∀*p*_*i*_ ∈ ^*•*^*t*_*j*_, |**M**_*k™*1_(*p*_*i*_)| ≥ 1); 2. *Predictive Safety:* The prognostic gating function verifies that the forecasted clinical trajectory satisfies an informatics-defined safety threshold (i.e., Φ(*t*_*j*_, **M**_*k™*1_) = 1).

If the automated validation fails, whether attempting to advance to surgical resection without required functional mapping, or prescribing an intervention computationally predicted to trigger a severe adverse event, the transition is instantly blocked. This rigorous formalization actively prevents diagnostic omissions, filters out unstructured noise, enforces evidence-based protocol adherence.

### Color semantics and dashboard realization

The interactive color coding that clinicians observe on the deployed dashboards is not a cosmetic display layer but the direct visual rendering of the colored marking. We make this correspondence precise. The status lattice is the finite chain Γ =( {∅, B, Y, G }, ⪯), ∅≺ B ≺Y≺ G, where ∅ denotes a step not yet initiated and B (Blue), Y (Yellow), and G (Green) denote, respectively, a diagnostic step that has been *ordered, scheduled*, and *completed*, exactly the indicator semantics of the Patient Journey Tracker (Sec. 4.4, Fig. 5). What the screenshots display is then the *status projection* of the live marking, an observation map *π*_Γ_ : *M* →Γ^*P*^ defined cell-wise by the join over the colored multiset resident at each place. Each color-coded cell in the tracker (Fig. 5) and each status badge in the EEG and EMU interfaces (Figs. 3–4) is therefore *literally π*_Γ_(**M**_*k*_)(*p*_*i*_): the combinatorial colored dashboard is a faithful, real-time isomorphic view of the underlying Petri-net state rather than an independently maintained artifact.

**Figure 3.**
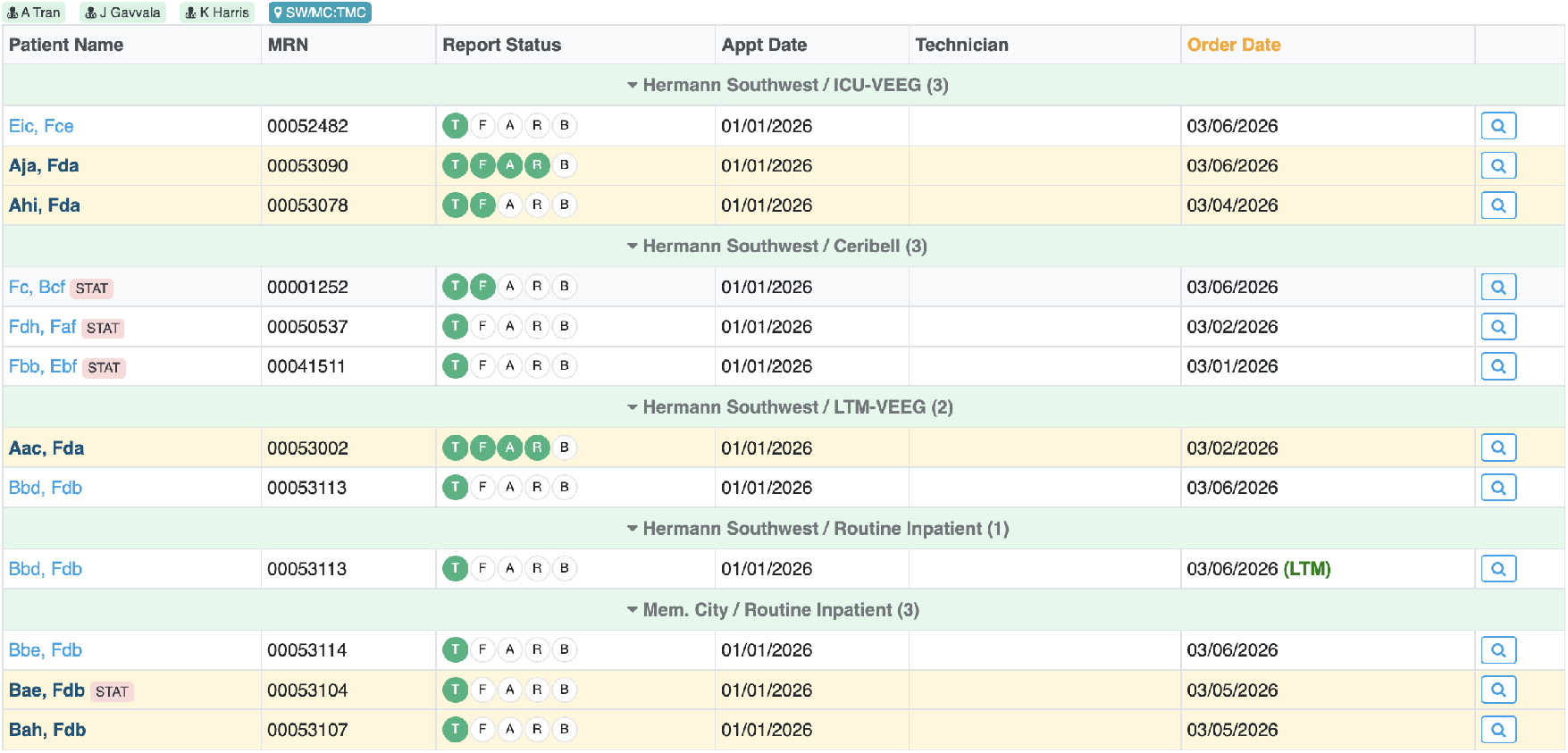
EEG report interpretation interface that allows physicians to document EEG findings using structured elements and automatically generate standardized clinical reports.

**Figure 4.**
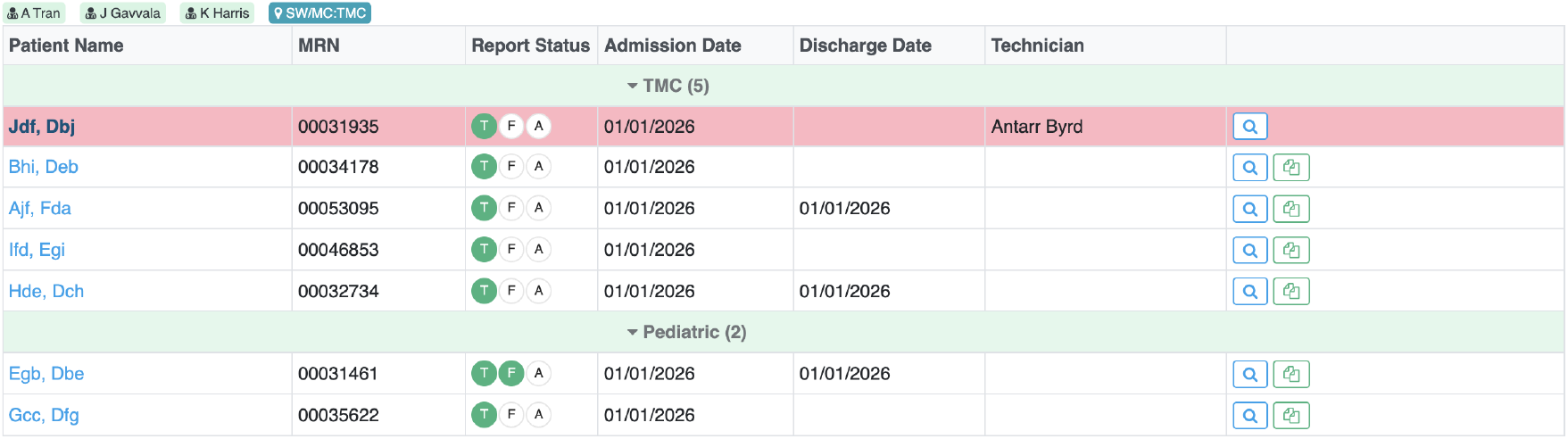
EMU phase reporting interface used to document seizure events and summarize monitoring results during inpatient epilepsy evaluation.

**Figure 5.**
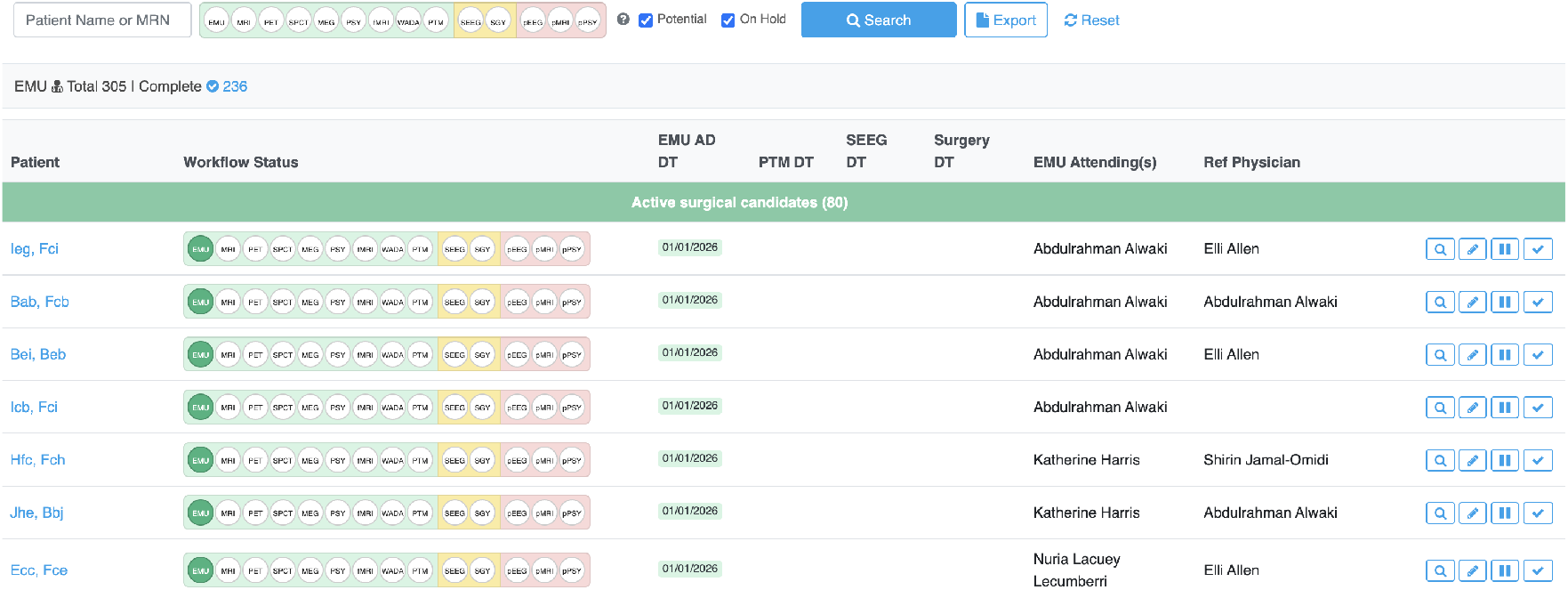
Patient Journey Tracker reflecting a 14-step surgical evaluation workflow using color-coded status indicators.

**Figure 6.**
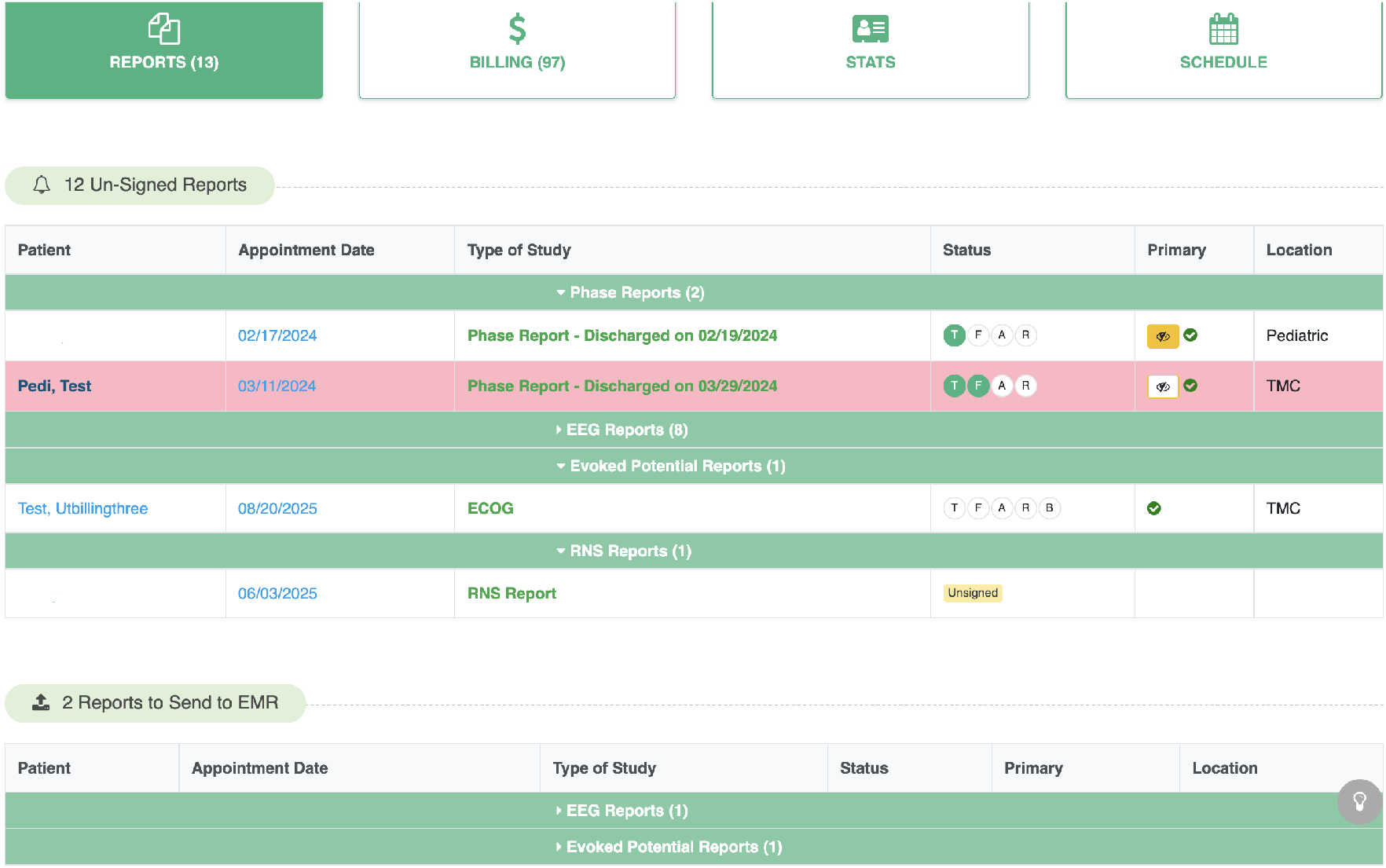
Physician dashboard integrating reporting, billing, statistics, and scheduling functions.

### Guiding Principles

We formulated the **PRSIVILL** principles that help guide EHT implementation.

#### Physician-Driven Design (P)

The system architecture, user interface, and data hierarchies of the EHT must mirror the clinician’s natural cognitive mental model. Development must employ agile, interface-driven methodologies involving attending physicians, rather than being dictated by hospital coders.

#### Reduced Clicks and Cognitive Load (R)

EHTs must aggressively automate repetitive data retrieval. The goal is to maximize the physician’s cognitive bandwidth for complex medical decision-making by pushing context-aware data to a unified dashboard.

#### Specialty-Specific (S)

Medicine is fundamentally heterogeneous. EHTs require high-resolution modeling of specific disease pathways (e.g., epileptology, neuro-oncology) rather than relying on generic, “one-size-fits-all” textual templates.

#### Interoperability (I)

EHTs must not create isolated data silos. They should be engineered as “tethered” systems that seamlessly synchronize with the institutional parent EHR (e.g., Epic) utilizing robust interoperability standards including HL7v2 and SMART on FHIR.

#### Value-Oriented (V)

Implementations must yield demonstrable, quantifiable benefits across three pillars: clinical outcomes (patient safety), care velocity, and administrative efficiency (throughput optimization).

#### Innovation Stack (I)

EHTs must leverage modern, scalable technological stacks such as cloud-native microservices, reactive web frameworks, and NoSQL document databases in order to support rapid deployment cycles.

#### Learning-Ready (L)

Following the vision of a continuous LHS, EHTs should capture highly structured, ontologically anchored data natively at the point of care, adhering to FAIR data principles^10^, and rendering clinical records ready for secondary use.

#### Longitudinality (L)

The twin should track the entire patient’s clinical care trajectory continuously across time, disparate care settings, and life stages, providing an uninterrupted temporal avatar of disease progression.

### 3.1 System Design and Implementation: The 14-Step EMU Digital Twin

The EHT framework has been embodied in the *Epilepsy Tracking optimized Management engine (EpiToMe)*^7^. Utilizing the Epilepsy and Seizure Ontology (EpSO^9^) as a foundational backbone, we mapped the pre-surgical evaluation workflow into a strict 14-step computational state machine. This workflow encompasses the entirety of the clinical trajectory: *EMU Admission, Scalp vEEG, 3T/7T MRI, FDG-PET, SPECT, MEG, fMRI, Wada Testing, Neuropsychology, Multidisciplinary Case Conference, Phase II Monitoring, Cortical Mapping, Surgery, and Post-Operative Follow-up*.

EpiToMe is deployed as a bespoke web-based application interfacing securely with the institutional enterprise EHR via an asynchronous, event-driven architecture (**Fig. 2**). When a clinical event occurs (e.g., a signed radiology report), a listener engine parses the HL7 broadcast in real-time, extracts clinical impressions using a hybrid Natural Language Processing (NLP) pipeline, maps terminology to EpSO, and autonomously transitions the patient’s digital twin state.

## 4 Results

### System Deployment

The EpiToMe EHT instance has been continuously deployed and expanded in the Epilepsy Center since February 2019. The system operates as a web-based clinical specialty digital twin integrated with the institutional enterprise Electronic Health Records through HL7 messaging interfaces and SMART on FHIR APIs. Clinical events such as EEG studies, EMU admissions, neuroimaging reports, neuromodulation-related documentation, and multidisciplinary case conferences are synchronized with the EHT to support structured documentation, longitudinal data capture, and workflow tracking.

Since its initial production deployment, EpiToMe has supported multiple epilepsy care workflows and has been routinely used by epileptologists, fellows, technologists, nurse navigators, and multidisciplinary care teams. As of today, EpiToMe has documented a total of 184,107 clinical reports across seven reporting interfaces for 41,719 patients (**Table 1**).

**Table 1.** EpiToMe Reports.

| Type | Production Since | Reports | Patients |
| --- | --- | --- | --- |
| EEG | 02/18/2019 | 165,248 | 36,544 |
| EMU Phase | 07/01/2019 | 7,140 | 2,290 |
| EMU Daily | 11/15/2019 | 10,869 | 2,162 |
| Evoked Potentials | 01/14/2021 | 260 | 250 |
| MEG | 08/16/2023 | 303 | 277 |
| Neuropace | 06/03/2025 | 143 | 78 |
| Care Conference | 02/07/2020 | 144 | 118 |
| Total |  | 184,107 | 41,719 |

Seven report authoring interfaces have been implemented within the system: *EEG report interpretation, EMU phase reports, EMU daily reports, care conference reports, evoked potentials reports, MEG reports, and Neuropace reports*.

EEG reporting was the first workflow deployed into production in February 2019, followed by EMU phase reporting in July 2019, EMU daily reporting in November 2019, care conference reporting in February 2020, evoked potentials reporting in January 2021, MEG reporting in August 2023, and Neuropace reporting in June 2025 (**Table 1**). EEG reporting represents the dominant production workload, with 165,248 reports generated for 36,544 patients. EMU workflows also account for substantial clinical use, including 7,140 EMU phase reports and 10,869 EMU daily reports, each supporting 2,162 patients. Collectively, these data demonstrate that the EHT framework supports both high-volume routine documentation and lower-volume specialized reporting within a digital twin production system.

EpiToMe is an empirical instantiation of the Colored Workflow Petri Nets model. The seven reporting interfaces realize seven distinct semantic color domains (EEG, EMU-phase, EMU-daily, care-conference, evoked-potential, MEG, and Neuropace), and the platform maintains one colored marking per patient. Every signed report is the observable trace of a status-promoting control event **u**_*k*_(*D*_in_) that advanced its token to the Green (G, Completed) status of Γ; the 184,107 reports therefore correspond to 184,107 *realized transition firings* across the twin model. Because each firing is admitted only under the semantic-conformance and predictive-safety guards, the cumulative marking constitutes the system’s empirically reachable state set: the structured, ontology-anchored substrate that EpiToMe’s cohort-discovery engine interrogates, and that the FAIR-compliant JSON export serializes without retrospective chart-review in the parent EHR.

The staged production timeline in Table 1 further reflects a *monotone enlargement* of the net: each newly deployed interface (from EEG reporting in 2019 through Neuropace reporting in 2025) adjoins additional places and a color set *τ* to Σ without invalidating the conformance invariants already verified on the existing sub-nets. This compositional extensibility is what allowed the workflow scope to expand by seven interfaces over six years while preserving a single, internally consistent state-machine semantics.

### EEG Report Interpretation Workflow

The EEG Report Interpretation Workflow represents a digital twin of the routine EEG reporting process. This workflow provides a structured interface that allows physicians to interpret EEG recordings and document findings using standardized clinical elements.

Within the reporting interface, clinicians can capture structured information regarding background rhythms, epileptiform discharges, seizure patterns, and clinical correlations. These interpretation elements are organized into structured domains that guide physicians through the reporting process while ensuring consistency of clinical documentation.

Once interpretation fields are completed, the system automatically generates a formatted EEG report that can be transmitted back to the parent EHR system with a single action. By structuring EEG interpretation data into standardized fields, the workflow reduces documentation complexity while enabling downstream computational analysis and research use.

### Epilepsy Monitoring Unit Phase Report Workflow

The Epilepsy Monitoring Unit (EMU) Phase Report Workflow models the inpatient presurgical epilepsy monitoring process as a digital twin of the EMU clinical evaluation pathway.

During EMU admission, patients undergo continuous video-electroencephalography monitoring to capture seizure events and characterize seizure semiology. The EHT system provides a structured interface that allows clinicians to document seizure characteristics, electrographic findings, medication adjustments, and clinical interpretations during the monitoring period.

The workflow organizes EMU documentation into structured reporting elements, enabling clinicians to efficiently summarize seizure activity and diagnostic findings during EMU evaluation. Structured documentation also enables automatic aggregation of seizure statistics and facilitates rapid generation of EMU phase reports.

### Patient Journey Tracker Workflow

Epilepsy care is a complex multidisciplinary process that involves neurologists, radiologists, neuropsychologists, and neurosurgeons, particularly for patients who are candidates for epilepsy surgery. To support coordination among these clinical teams, EpiToMe implements a Patient Journey Tracker that functions as a longitudinal digital twin of the patient’s clinical pathway.

The tracker records fourteen key steps in the epilepsy surgical evaluation process, including EMU admission, EMU discharge, PET imaging, ictal SPECT, MEG, neuropsychology testing, functional MRI, WADA testing, patient management conferences, SEEG implantation, surgical intervention, and post-operative follow-up evaluations including EEG, MRI, and Neuropsychology.

Color-coded indicators are used to represent the status of each step within the workflow. Blue indicates that a diagnostic test has been ordered, yellow represents a scheduled procedure, and green indicates that a step has been completed. This visual representation allows clinicians to quickly identify pending evaluations and workflow bottlenecks.

The tracker also provides advanced search functionality that allows clinicians to identify patient cohorts based on workflow status. In addition to searching by patient name or medical record number, users can search patients by completion status of specific diagnostic steps and combine multiple criteria to identify relevant cohorts.

### Physician Dashboard

To support physician-centered workflow design, EpiToMe provides an integrated Physician Dashboard that serves as the primary interface for daily clinical activities. When physicians log into the system, they are automatically directed to the dashboard corresponding to the current date, allowing them to manage clinical tasks without navigating across multiple systems.

The dashboard consists of four primary functional tabs: Reports, Billing, Statistics, and Schedule.

The *Reports* tab displays all outstanding reports requiring physician review or completion. Physicians can finalize reports within the system and transmit completed reports back to the parent EHR with a single action.

The *Billing* tab automatically retrieves billing-related information associated with completed reports and presents them in a structured interface. Physicians can submit billing documentation with only three to four clicks. In contrast, completing the same billing process in the enterprise EHR system typically requires more than twenty clicks across multiple pages.

The *Statistics* tab provides an overview of reporting activities, including the number of reports completed per month and the distribution of reports by study type.

The *Schedule* tab allows physicians to review their service schedules throughout the year and request schedule exchanges with other physicians. The schedule functionality also enables EpiToMe to automatically associate reports with the responsible physicians.

### Surgical Planning Tool for Care Conference

To support the specialized pre-surgical evaluation workflow, the EHT framework incorporates a dedicated Surgical Planning Tool^13^. This module serves as a collaborative interface for SEEG and Subdural Electrode (SDE) strategic planning during multidisciplinary care conferences.

Traditional SEEG planning is often hindered by a “visual rendering” gap, where clinical teams are often forced to choose between simple hand-drawings that lack detail, or complex surgical software that lacks interoperability. The tool overcomes these limitations by implementing a neuroanatomical rendering engine that provides six interactive, ontology-anchored views of a template brain atlas (Figure 7).

**Figure 7.**
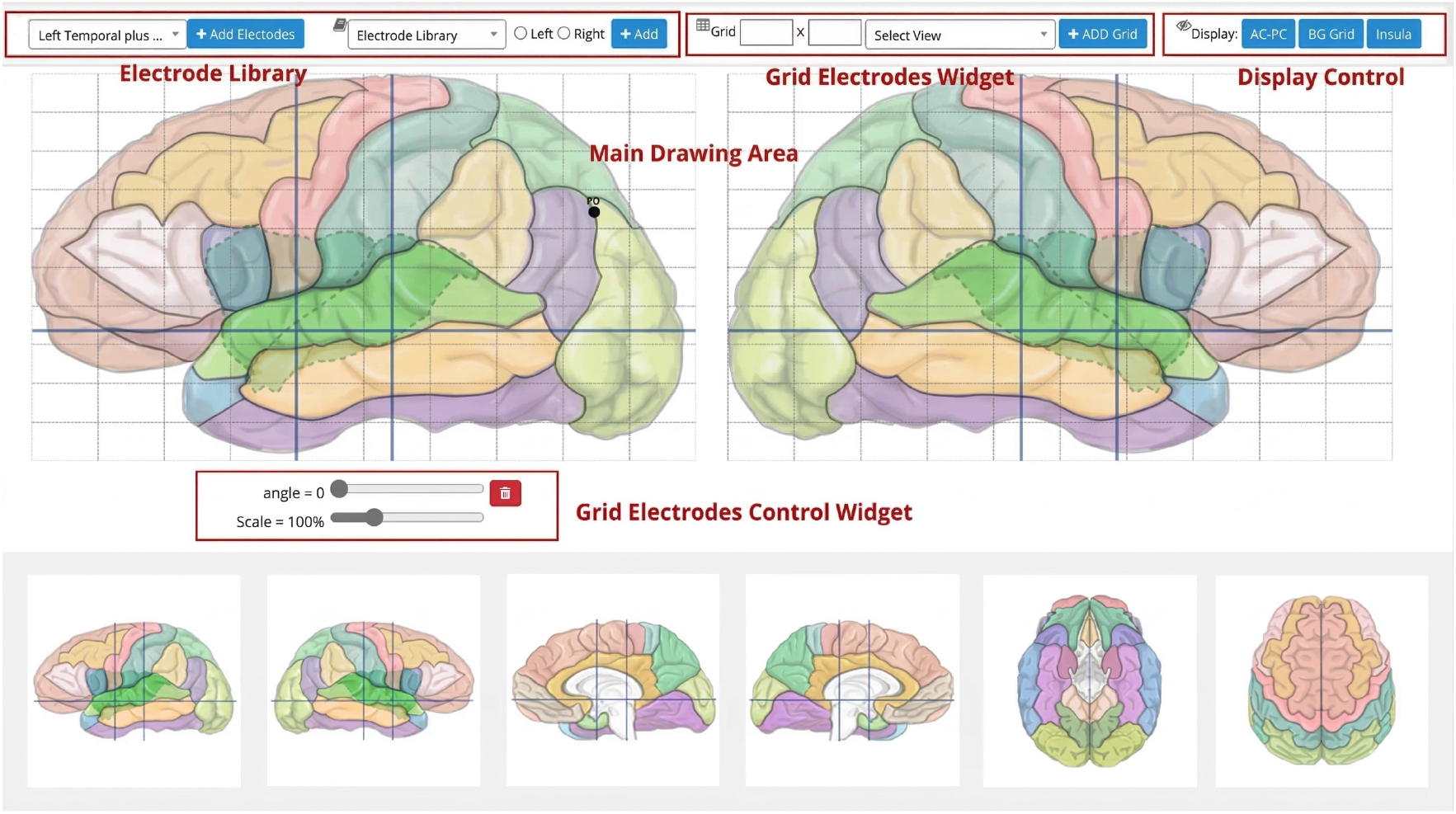
The Surgical Planning Tool interface featuring a six-view neuroanatomical atlas. The tool integrates a standardized electrode library to ensure consistent terminology and collaborative documentation of electrode trajectories.

Central to the tool’s collaborative efficiency is the *standardized electrode and electrode group Common Data Elements*, which centralizes institutional expertise into a knowledge repository. This library features 52 predefined electrode types and 14 conventional group configurations, ensuring that all multidisciplinary team members adhere to consistent clinical terminology and anatomical targets. By centralizing these best practices, the tool eliminates naming ambiguity during high-stakes planning sessions and functions as a vital educational platform. Trainees, such as residents and fellows, can utilize this library to familiarize themselves with established clinical standards and implantation strategies, effectively shortening the learning curve for complex surgical planning.

The tool ensures that clinical intent is captured as structured, FAIR-compliant data. In a longitudinal evaluation (2022–2024), this tool captured over 6,600 electrode trajectories with a System Usability Scale (SUS) score of 80.4, effectively bridging the gap between clinical discussion and structured documentation.

### Beyond Evaluation: Impact Assessment

#### Quantitative Impact on Clinical Productivity

The implementation of the EHT demonstrated statistically significant improvements in operational efficiency. Under the legacy EHR model, the average time required for a specialist to manually aggregate multimodal diagnostic data across disparate tabs to synthesize a Phase I Case Conference report was 55.4 (± 12.3) minutes per patient. Following EHT deployment, automated data synthesis reduced this duration to 14.2 (± 4.1) minutes, representing a 74.4% reduction in documentation burden (*p <* 0.001). By alleviating documentation bottlenecks like this, the epilepsy center sustained a nearly 400% increase in the annualized volume of completed surgical evaluations without necessitating an increase in attending physician or support staff headcount.

#### Informatics Paradigm Shift: Structured Data Fidelity

A foundational outcome of the EHT framework was catalyzing the structural transition from document-centric to data-centric medical informatics (Table 2). Because the state machine mathematically forces clinical inputs into specific pathways, the system achieved near complete capture of specialized clinical phenotypes as highly structured, ontologically mapped fields. Consequently, longitudinal patient records were natively exported as FAIR-compliant JavaScript Object Notation (JSON) structures, eliminating the need for costly retrospective manual chart abstraction for clinical research.

**Table 2.** Paradigm Feature Comparison: EHR vs. EHT (EpiToMe)

| Informatics Dimension | Legacy Monolithic EHR | EHT Framework (EpiToMe) |
| --- | --- | --- |
| Data Entry Mode | Narrative free-text notes, disjointed tabs | Structured, ontology-mapped discrete fields |
| State Tracking | Manual chart biopsy / mental tracking | 14-step Petri-Net state machine |
| Cognitive Paradigm | System of Record (Billing optimization) | System of Care (Care pathway optimization) |
| Research Readiness | Burdensome retrospective abstraction | FAIR-compliant, analytic-ready JSON |

#### Qualitative Physician Feedback and Patient Safety

In prior formal qualitative evaluations of clinical users (*N* = 25), 100% of respondents affirmed that the structured, visual 14-step EHT dashboard improved perceived patient safety.^7^ The strict mathematical enforcement of diagnostic prerequisites via the WPN architecture effectively prevented the omission of critical preoperative tests during complex neurosurgical planning, thereby directly mitigating factors contributing to clinical cognitive overload.

## 5 Discussion

The empirical success of the EpiToMe system presents a highly scalable model for specialized medical informatics. By abstracting complex clinical logic into a bespoke application tethered to the enterprise EHR via SMART on FHIR APIs, organizations can deliver specialized cognitive tools without fragmenting the institutional data backbone.

### Bridging Enterprise and Medical Digital Twins

Beyond operational efficiency, the EHT framework directly addresses the translational “data silo” problem. Because operational phenotypes are natively captured as highly structured data, they serve as optimal clinical anchors for multi-scale biological research. To demonstrate ultimate scalability, the dataset generated by EpiToMe currently functions as the clinical foundation for the “Tissue-to-Bytes” consortium initiative.^11^ Integrating deep single-cell omics, spatial transcriptomics, and high-resolution digital pathology directly into the patient’s operational twin establishes a multi-scale digital twin spanning macroscopic clinical behavior to the molecular architecture of resected tissue. This paves the way for a true Learning Healthcare System, wherein consortium-derived transcriptomic insights can be seamlessly codified into the EHT’s feed-backward decision support logic at the point of care.

### Limitations and Future Work

While the findings demonstrate substantial informatics advancements, several limitations must be acknowledged. First, the evaluation relies on a single-center implementation within a highly specialized tertiary academic domain (epileptology). Generalizability to community hospital settings with differing IT infrastructures or to other medical specialties (e.g., transplant cardiology) requires multi-center validation. Second, the EHT’s real-time accuracy remains strictly dependent on the latency and quality of unstructured data originating from the parent EHR via HL7 feeds, necessitating periodic manual clinician overrides. Third, maintaining complex domain ontologies (such as EpSO) requires ongoing, dedicated informatics governance to prevent conceptual drift. Future work will focus on integrating predictive machine learning models into the feed-backward loops. In doing so, stringent ethical guardrails, such as the mandatory use of Clinical Model Cards^12^ to document algorithmic biases, will be essential to ensure that the EHT strictly augments, rather than replaces, human clinical judgment.

## 6 Conclusion

Conventional monolithic EHR architectures predominantly function as administrative repositories, often lacking the dynamic capabilities necessary to optimally facilitate patient care. The Enterprise Health Twin framework presents a theory-grounded methodology that integrates the cybernetic principles of Digital Twin technology into clinical care workflows. Based on a seven-year longitudinal deployment of the EpiToMe system, our findings empirically demonstrate that modeling clinical processes under the EHT framework significantly reduces provider cognitive load, enhances operational throughput, and natively generates FAIR-compliant clinical data at the point of care. By emphasizing interoperable standards and facilitating the linkage between operational workflows and patient care journeys, this framework provides a viable pathway for learning health systems and beyond^17,18^.

## Acknowledgment

This work was supported by the National Science Foundation (NSF) through grant IIS2500624, and National Institutes of Health (NIH) through grants R24NS143946, R01NS126690, R01NS116287, U24MH130988, and U24AG098157. The content is solely the responsibility of the authors and does not necessarily represent the official views of the NSF or NIH.

